# Investigating the effect of mechanical adaptation on mid-air ultrasound vibrotactile stimuli

**DOI:** 10.1101/2024.12.11.627964

**Authors:** Antonio Cataldo, Tianhui Huang, William Frier, Patrick Haggard

## Abstract

Gesture control systems based on mid-air haptics are increasingly used in infotainment systems in cars, where they can provide rich haptic feedback to improve human-computer interactions. Laboratory studies show that mid-air haptic feedback reduces drivers’ distractions and improve safety. However, it is unclear how the perception of mid-air ultrasound stimuli is affected by prolonged exposure to vibrational noise, e.g., from the steering wheel of a moving vehicle. Studies on vibrotactile adaptation show that perception of mechanical vibration is impaired by prior exposure to stimuli of the same frequency. Here, we investigated the effect of mechanical adaptation on the perception of mid-air ultrasound stimuli. We measured participants’ detection threshold for ultrasound stimuli of different frequencies both before and after exposure to 30 s mechanical vibrations. Across two experiments, we systematically manipulated the frequency and amplitude of the adapting stimulus. We found that exposure to low-frequency mechanical vibrations significantly impaired the detection of low-frequency ultrasound stimuli. In contrast, exposure to high-frequency mechanical vibrations equally impaired perception of both low- and high-frequency ultrasound stimuli. This effect was mediated by the amplitude of the adapting stimulus, with stronger mechanical vibrations producing a larger increase in participants’ detection threshold. Overall, these findings show that perception of mid-air ultrasound stimuli is affected by specific sources of mechanical noise. Crucially, frequency-specificity in the low-frequency band also points toward possible mitigating solutions that could help minimising unwanted desensitization of mechanoreceptor channels during mid-air haptic interactions.

## I. INTRODUCTION

Mid-air ultrasound stimulation has the potential to revolutionise haptic technologies in many different fields, including virtual reality, telecommunication, gaming, and Human-Computer Interactions in general [1], [2], [3], [4]. In the context of the automotive industry, a fast-growing body of laboratory studies has shown that ultrasound haptic feedback can enhance the driver’s sense of control over gesture-based interactions with the vehicle infotainment system, reducing drivers’ distractions and improving safety [5], [6], [7], [8], [9], [10], [11], [12].

However, it remains unclear whether and how the perception of mid-air haptic stimuli is affected by prolonged exposure to mechanical vibrations. Addressing this question has important implications for the use of mid-air haptics in environments that are subject to different sources of vibrational noise [12], [13]. For example, in the context of mid-air gesture-control systems in cars, the driver’s hand is likely to be exposed to protracted mechanical vibrations from the steering wheel prior to interacting with the infotainment system. Therefore, quantifying any effect of mechanical noise on the user’s sensitivity to mid-air haptic feedback is paramount for the safe implementation of mid-air gesture-based interactions in real world scenarios.

Vibrotactile adaptation is a well-known phenomenon in the somatosensory literature, whereby exposure to protracted mechanical vibrations induces a decrease in the perception of subsequent vibrotactile stimuli delivered to the same stimulation site [14], [15], [16], [17], [18], [19], [20], [21], [22]. Previous studies have shown that adapting stimuli of either low- or high-frequency selectively impair detection of test stimuli of similar frequency only [14], [15], [23], [24], [25], [26], but see [27], [28]. This frequency-specific effect is commonly explained as the result of fatigue of the sensory receptors within a specific mechanoreceptive channel when both the adapting and the test stimuli have similar frequency. Similarly, the lack of adaptation between stimuli with different frequency is generally thought to reflect the preferential activation of distinct mechanoreceptive channels [28], [29], [30], [31], [32], [33].

Crucially, virtually all the previous studies on vibrotactile adaptation have used mechanical stimulation both as adapting and test stimulus. However, the human glabrous skin is innervated by three main types of cutaneous receptors, each preferentially responding to specific mechanical deformation of the skin [34], [35]. The slow adapting type I channel (SA-I) innervates Merkel receptors and is most sensitive to static pressure. The rapidly adapting (RA) channel innervates Meissner receptors and is preferentially activated by low-frequency, flutter-like vibrations (10-50 Hz). The Pacinian channel (PC) takes its name from the Pacinian corpuscle it innervates, and is mostly sensitive to high-frequency vibrations (> 50Hz, peaking at 250 Hz).

The use of mechanical stimulation in previous studies means that a constant indentation must be applied to the participant’s skin in order to deliver the vibrotactile stimuli. As a consequence, previous results may at least partially reflect interactions between mechanical propagation waves on the skin [36], [37]. In contrast, mid-air ultrasound stimulation does not require mechanical contact with the skin, and therefore it has the potential to provide purely frequency-resolved stimuli. This is supported by preliminary microneurography data suggesting that mid-air ultrasound stimuli do not activate SA receptors [38]. Therefore, it remains unclear how a mechanical adapting stimulus might affect the perception a target mid-air ultrasound stimulus.

To our knowledge, only one previous study to date has investigated the interaction between mechanical noise and ultrasound perception [39]. In that study, Špakov et al. compared the ability of a driver to recognise the shape of four different mid-air stimuli both while driving and when vibrational noise was absent. Although they found that vibrational noise had little effect on the participants’ ability to recognize the shape of the mid-air stimulation, their ecological study did not systematically manipulate the frequency and amplitude of neither the mechanical nor the ultrasound stimulation, and therefore was unable to investigate the effect of vibrotactile adaptation. Thus, it remains unclear whether the findings from vibrotactile adaptation for mechanical stimuli also apply to ultrasound stimuli, and how the perception of mid-air ultrasound stimulation might be affected by prolonged exposure to mechanical vibrations.

Here, we directly address this question in two different experiments based on power calculation, replication, and preregistration. In Experiment 1, we measured participants’ detection threshold for mid-air ultrasound stimulation both before and after adaptation to a 30 s mechanical vibration. Mechanical adapting vibrations were delivered by means of a robotic arm with independent control of both frequency and amplitude, while mid-air stimuli were delivered using an ultrasound array to project acoustic pressure points on the hand with high levels of spatiotemporal precision. To investigate the frequency-specificity of adaptation, we systematically manipulated the frequency of both the adapting mechanical stimulus, and the test ultrasound stimulus. Based on the classical observation of frequency-specific vibrotactile adaptation [14], [15], [23], [24], [25], [26], we hypothesised and preregistered (see https://osf.io/zwmnt) a significant interaction between the specific frequency of mechanical and ultrasound stimulation, whereby a low-frequency mechanical adapting stimulus would selectively impair the detection of a low- but not high-frequency ultrasound test stimulus, and vice versa.

Previous studies on vibrotactile adaptation have shown a linear relationship between the amplitude of the adapting stimulus and the increase in detection threshold [20], [21], [40], [41]. Therefore, besides providing a direct replication of our first experiment, Experiment 2 investigated whether the detectability of a test ultrasound stimulus was affected by the amplitude of the mechanical adapting stimulus. Based on the literature cited above, we hypothesised and preregistered (see https://osf.io/g8jft) that stronger adapting stimuli would produce a significantly higher increase in detection threshold of ultrasound test stimuli.

## II. EXPERIMENT 1

### A. METHODS

#### 1) PARTICIPANTS

The final sample size (n = 12; 10 females; mean age ± SD: 23.4 ± 2.8) was determined a-priori through a power analysis, based on the results of a pilot study with a similar design involving six participants (see preregistration at https://osf.io/zwmnt for details). With an alpha level of 0.05 and a power of 0.80, the estimated sample size required to demonstrate a significant difference in detection thresholds was twelve participants.

A total of 15 right-handed healthy volunteers were originally recruited for the study. Based on preregistered exclusion criteria (https://osf.io/zwmnt), three participants were excluded after visual inspection of the data because they showed poor convergence (i.e., > 25% discrepancy) between ascending and descending staircases in the main task (see Procedure).

All the methods and procedures employed in the study adhered to the Declaration of Helsinki and were approved by the UCL Research Ethics Committee. All participants were naïve regarding the hypotheses of the study, provided their written informed consent before the beginning of the experiment, and received a cash compensation of £9 per hour.

#### 2) APPARATUS AND STIMULI

The experimental setup is depicted in Figure 1A-C. Participants sat in front of a desk, resting their left elbow on an articulated armrest (YANGHX, model 4328350928, China). Their left hand was placed palm down on a plastic support, in correspondence of a 10 cm × 10 cm opening. Mid-air ultrasound stimulation (i.e., test stimulus) was delivered via a STRATOS Explore device (Ultraleap, UK) located 12 cm below the participant’s hand (Figure 1A-B). The STRATOS Explore is a computer-controlled device comprising a grid of 16 × 16 transducers that utilizes focused ultrasound (40 KHz) to project discrete tactile points directly onto the participants’ skin [1], [4], [42]. This device allowed us to design mid-air stimulations of different frequencies and amplitudes. In particular, the test stimulus consisted of a single focal point (6 mm diameter) delivered for 1 s to the centre of the participant’s palm of the left hand.

**FIGURE 1.**
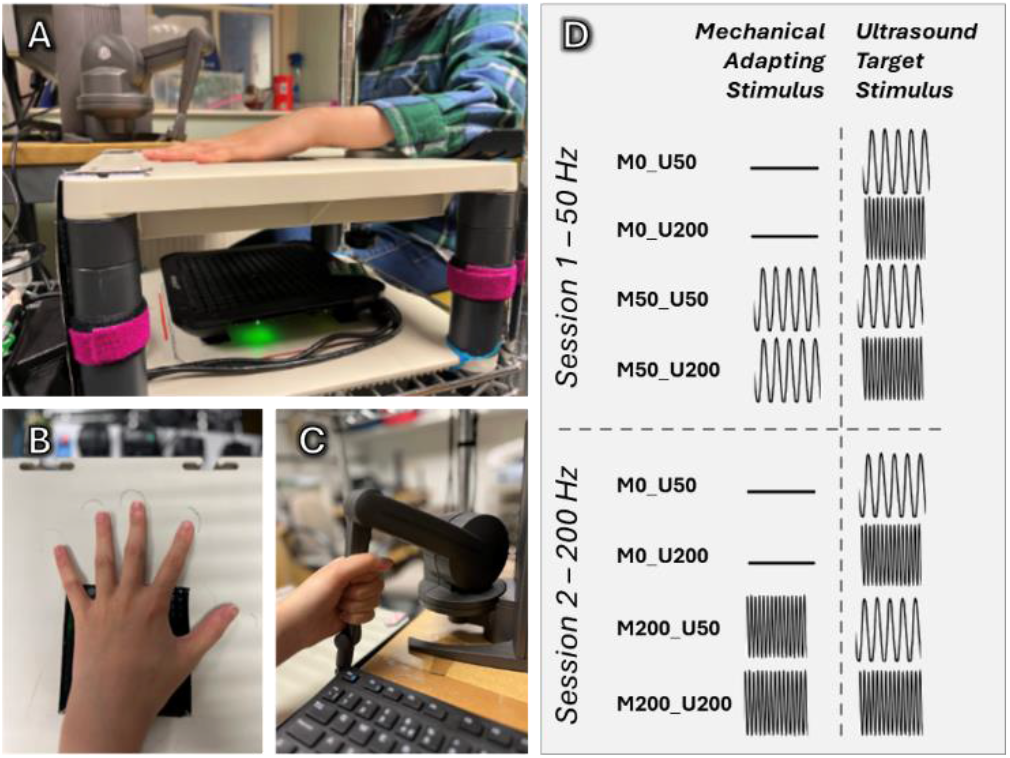
Experimental Setup. **A**. Participants sat at a desk and rested their left hand on a plastic support, in correspondence of a 10 × 10 cm opening. **B**. Mid-air ultrasound stimulation (i.e., target stimulus) was delivered via a STRATOS Explore device located 12 cm below the participant’s hand. **C**. During the mechanical stimulation (i.e., adapting stimulus), participants were asked to hold the robotic arm of a Geomagic Touch X device with their left hand, and to ensure that the center of their palm was fully in contact with the device. **D**. The experiment took place in two separate sessions, one for each mechanical adaptation frequency. Within each session, participants performed four blocks, including a baseline, no-adaptation condition, and a post-adaptation condition for each ultrasound stimulus.

The amplitude of the test stimulus was modulated using a sinusoidal signal at either 50 Hz or 200 Hz in different conditions, to preferentially activate the low-frequency RA channel or the high-frequency PC channel, respectively [34], [43]. The amplitude of the mid-air stimuli ranged from 0 to the maximal output achievable with the STRATOS Explore device, and was adjusted adaptively in a psychophysical staircase procedure to establish the participant’s detection threshold (see Procedure).

The mechanical vibrotactile stimulation acting as adapting stimulus was delivered through a robotic arm (Geomagic, Touch X, USA). During the adaptation phase, participants were asked to hold the robotic arm with their left hand (see Figure 1C), such that the mechanical stimulation was delivered to the centre of their palm, at the same stimulation site of the ultrasound test stimulus. The adapting stimulus lasted for 30 s. The haptic force feedback capabilities of the Geomagic Touch X device allowed us to specify a time-varying force waveform such to independently manipulate the frequency and amplitude of the adapting mechanical stimulus.

To test the effect of frequency-specific adaptation, the frequency of the adapting stimulus was set at either 50 Hz (preferentially activating the RA channel) or 200 Hz (preferentially activating the PC channel) [34] in different conditions. The amplitude of the 200 Hz adapting stimulus was set to 1.5 N. This was based on the results of two pilot experiments with a similar design conducted before the formal experiment (pilot 1: n = 3, pilot 2: n = 6. These pilot studies showed that with our device, amplitudes above 1.5 N produced a ceiling effect, completely suppressing perception of subsequent ultrasound test stimuli. The amplitude of the 50 Hz adapting stimulus was instead adjusted for each participant such that it matched the individual perceived intensity of the 1.5 N-200 Hz stimulus (see Procedure). This was done because the perceived intensity of a vibrotactile stimulus is strongly affected by its frequency [34]. Thus, establishing the perceived isointensity between adapting stimuli of different frequency ensured that the effect of mechanical adaptation in our study was not confounded by differences in perceived intensity.

The physical peak-to-peak skin displacement induced by each mechanical stimulation condition was measured in a validation experiment using accelerometery (ADXL335 Module Analog Accelerometer, Sparkfun) and Laser Doppler Vibrometry (LDV PSV-500-Scanning-Vibrometer, Polytec).

The experiment also included a baseline condition where the participant held the robotic arm with their left hand for the same duration as in the other conditions, but the Geomagic Touch X device was switched off. This allowed us to measure the participant’s detection threshold for mid-air ultrasound stimuli in absence of adaptation, while at the same time keeping task demands and other extraneous variables constant across conditions (e.g., contact pressure with the robotic arm, duration of adaptation phase, etc).

#### 3) PROCEDURE

##### Intensity matching task for mechanical stimuli

Before the beginning of the experiment, participants performed an intensity matching task to establish the perceived iso-intensity of a mechanical 50 Hz vibration matched to a 200 Hz mechanical vibration. The intensity matching task was based on a 2-AFC adaptive staircase procedure testing participants’ point of subjective equality (PSE) or perceived iso-intensity levels for the two mechanical adapting frequencies (50, 200 Hz). Participants held the robotic arm of the Geomagic Touch X device with their left hand (Figure 1C) and received two consecutive 1-s vibrations, separated by an interstimulus interval of 500 ms. Each stimulus was associated with a beep. After the second beep, participants were asked to report which beep contained the strongest vibrotactile stimulus. The intertrial interval was 500 ms from the participant’s keypress. The first stimulus in the trial (i.e. reference stimulus) was a 200 Hz stimulus with an amplitude of 1.5 N. The second stimulus (i.e. comparison stimulus) was a 50 Hz stimulus whose intensity was adaptively adjusted according to the participants’ response to the previous trial. A response indicating that the reference stimulus was felt as stronger produced an increase in the intensity of the comparison stimulus in the next trial. Conversely, a response indicating that the comparison stimulus was felt as stronger than the reference produced a decrease of the intensity of the comparison stimulus in the next trial. Two randomly interleaved ascending (starting from 0 N) and descending (starting from 0.75 N) staircases were used to test convergence and prevent response biases [44]. Increasingly smaller step-sizes (0.2, 0.05, 0.02 N) were used after the first two reversals, to provide a fast convergence toward the psychophysical thresholds. The maximal amplitude for the descending staircase and the amplitude of the step-sizes was determined in two previous pilot experiments (pilot 1: n = 3, pilot 2: n = 6). Each staircase ended after ten reversals. The average of the last eight reversals was used as an estimate of the perceived intensity ratio between the 50 Hz and the 200 Hz stimulus.

##### Detection threshold task for ultrasound stimuli

The main phase of the experiment had a 3 (mechanical adapting stimulus: no adaptation [baseline], 50 Hz, 200 Hz) x 2 (ultrasound test stimulus: 50 Hz, 200 Hz) within-participants factorial design. The 50 Hz and 200 Hz mechanical adaptation conditions were tested in two sessions taking place on separate days (Figure 1D). This was done to ensure complete wash out of any after-effects produced by a specific mechanical frequency and avoid unintended interference between the two mechanical stimulation conditions. To limit the effect of potential confounding factors affecting vibrotactile perception (e.g., temperature and humidity; [30]), the baseline detection thresholds for both the 50 Hz and 200 Hz ultrasound stimuli were tested at the beginning of each session. Thus, each session consisted of four blocks, two testing the baseline detection thresholds for both ultrasound test stimuli, and two testing the same detection thresholds after adaptation to a specific mechanical stimulation.

The order of the mechanical adapting stimulus conditions was counterbalanced across participants to control for any order effects, whereas the order of the ultrasound stimulation within the same mechanical adapting frequency was fully randomized.

Participants’ detection threshold for ultrasound test stimuli was established using a psychophysical staircase procedure similar to the one described above for intensity matching of mechanical stimulation. In each trial, participants received a single ultrasound test stimulus delivered to the centre of their left palm. The stimulus was accompanied by a beep, and participants were asked to report whether they felt any stimulation during the beep. The intensity of the ultrasound stimulus was adjusted depending on the participants’ response to the previous trial. A no detection response produced an increase in the intensity of the stimulus whereas a yes response produced a decrease in the intensity of the test stimulus. Two randomly interleaved staircases (ascending and descending), and three step sizes (20%, 10%, and 5% of the maximum output of the STRATOS Explore device) were again employed. Each staircase ended after 10 reversals, and the detection threshold was defined as the average of the last eight reversals.

At the beginning of each block, participants were asked to hold the robotic arm of the Geomagic Touch X device. After grasping the robotic arm, they received a 30 s mechanical adapting stimulation and were then asked to place their hand back on the plastic support. To ensure accurate timing of the adapting phase, the instructions were given through pre-recorded vocal messages, and participants were familiarised with the procedure prior to the beginning of the experiment. The adapting stimulation was repeated after every five trials of the staircase procedure.

Each block lasted approximately seven minutes, resulting in a testing session lasting around 70 minutes, including familiarization and instructions. Short breaks were provided between blocks and participants were instructed to move and stretch their bodies to avoid any postural discomfort from the prolonged stimulation.

##### Statistical Analyses

Participants’ detection thresholds for both ultrasound test conditions (50 Hz, 200 Hz) in each of the three mechanical adaptation conditions (baseline, 50 Hz, 200 Hz) were tested using a series of within-participants repeated measures ANOVAs, followed up by Bonferroni-corrected post-hoc analyses where appropriate. As from our preregistered analyses (https://osf.io/zwmnt), the raw data were used to compute the percentage change in detection threshold from to the baseline for each condition.

Non-significant results were further investigated with Bayesian factor analyses. Data analyses were performed using R48 and JASP v. 0.16.4 (JASP Team 2016, University of Amsterdam).

### B. RESULTS

The average perceived isointensity of the 50 Hz mechanical stimulus matched to the 200 Hz mechanical stimulus was (mean ± SD): 0.27 ± 0.15.

Regarding the absolute detection thresholds for ultrasound stimulation, we first performed a sanity check analysis to test whether the baseline thresholds were significantly different across the two testing sessions. A 2 (session: day 1, day 2) x 2 (ultrasound frequency: 50 Hz, 200 Hz) repeated measures ANOVA showed a significant main effect of ultrasound frequency 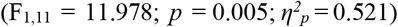, suggesting that the detection thresholds for the 200 Hz ultrasound stimulation (day 1: 0.36 ± 0.08; day 2: 0.35 ± 0.08) were significantly lower than the threshold for the 50 Hz stimulation (day 1: 0.50 ± 0.17; day 2: 0.52 ± 0.20), in line with previous results [43]. Crucially, the main effect of sessions was not statistically significant 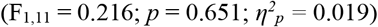. This null result was supported by a Bayesian factor analysis showing that the data were more than three times more likely under the null rather than alternative hypothesis (B_01_ = 3.138, error% = 0.021). Thus, the participants’ detection threshold for ultrasound stimuli was highly consistent across testing sessions, ruling out the possibility of any potential confounding factor intervening between sessions.

We therefore averaged together the baseline detection thresholds across the two testing sessions and ran a 3 (mechanical adapting stimulus: no adaptation, 50 Hz, 200 Hz) x 2 (ultrasound test stimulus: 50 Hz, 200 Hz) repeated measures ANOVA. The six conditions of this factorial design are described by the codes M0_U50, M0_U200, M50_U50, M50_U200, M200_U50, M200_U200. The results showed a significant main effect of ultrasound test stimulus 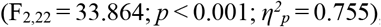, suggesting that participants’ detection thresholds were overall significantly higher for the 50 Hz test stimulus than the 200 Hz stimulus (see Figure 2A-B). Importantly, the main effect of mechanical adapting stimulus 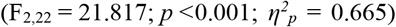 and the interaction between the two factors 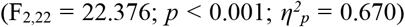 were also significant. To interpret the significant interaction, we ran a series of Bonferroni-corrected pairwise comparisons (see Table 1).

**Table 1:** Post-hoc pairwise comparisons of mean detection thresholds.

| Conditions | Mean diff.<br>± SD | t-value<br>(df = 11) | p-value* | Cohen's d |
| --- | --- | --- | --- | --- |
| M0_U50 vs M50_U50 | -0.25 ± 0.14 | -6.14 | 0.001 | 0.993 |
| M0_U50 vs M200_U50 | -0.15 ± 0.15 | -3.57 | 0.028 | -1.771 |
| M0_U200 vs M50_U200 | -0.04 ± 0.10 | -1.43 | 1.000 | -1.031 |
| M0_U200 vs M200_U200 | -0.18 ± 0.09 | -6.66 | < 0.001 | -0.413 |
| M0_U50 vs M0_U200 | 0.16 ± 0.16 | 3.44 | 0.042 | -1.921 |
| M50_U50 vs M50_U200 | 0.37 ± 0.11 | 11.56 | < 0.001 | 3.338 |
| M200_U50 vs M200_U200 | 0.13 ± 0.18 | 2.54 | 0.196 | 0.733 |

\*Bonferroni-corrected for seven comparisons.

**FIGURE 2.**
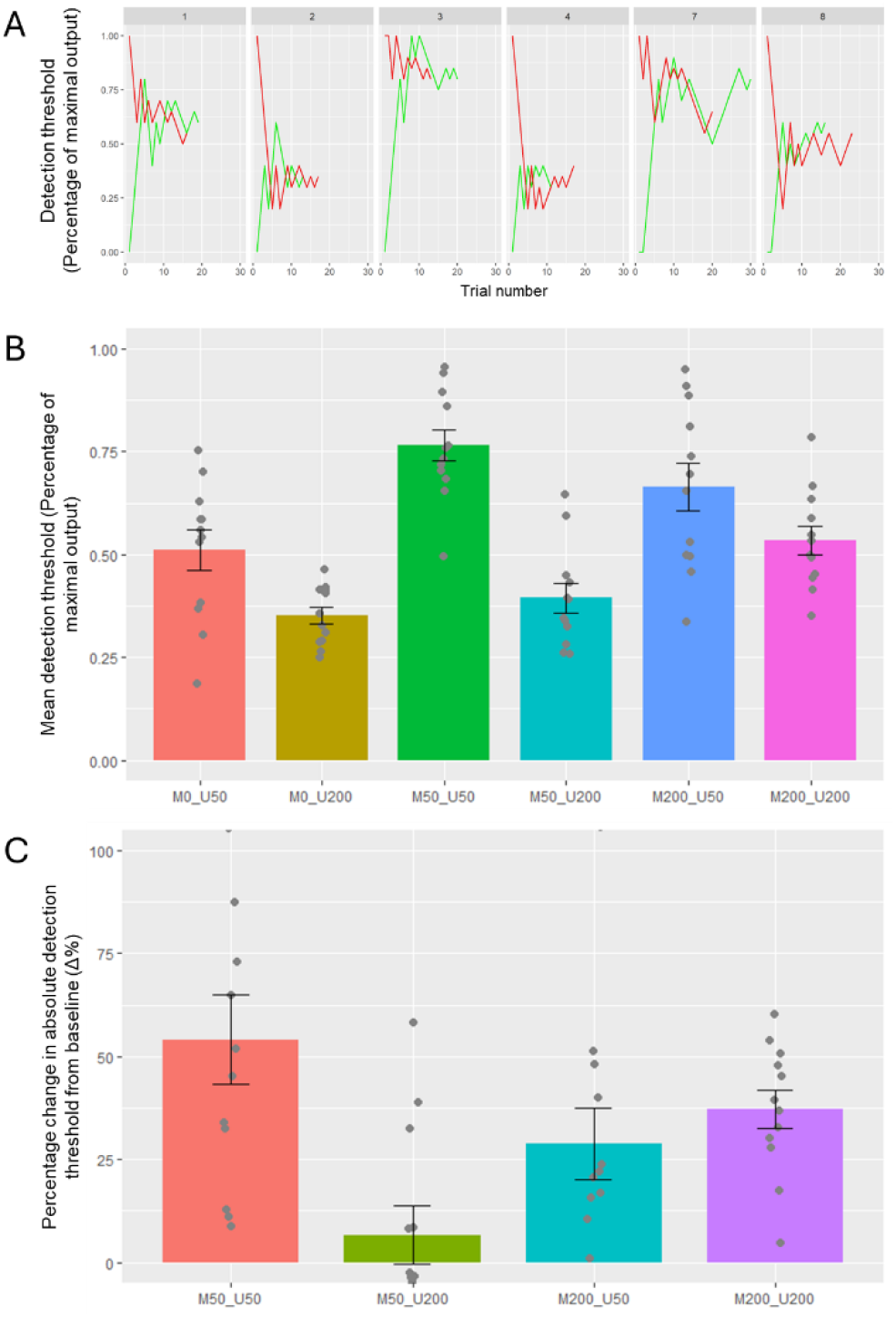
Results from Experiment 1. **A**. Staircase data from a representative participant (P# 14) across the different adaptation conditions (baseline data are shown for day 1 only). Green and red lines show responses to ascending and descending staircases, respectively. **B**. Mean absolute detection thresholds for 50 Hz and 200 Hz ultrasound stimuli in the three mechanical adaptation conditions (no adaptation, 50 Hz, and 200 Hz). The condition code MXX_UXX refers to the levels of the mechanical adapting stimulus (M) and the ultrasound test stimulus (U) respectively. **C**. Percentage change in absolute detection thresholds from baseline.

The analyses showed that all contrasts were significant (*p* ≤ 0.042 in all cases), except the difference between 200 Hz ultrasound thresholds before and after adaptation to 50 Hz mechanical stimulation, and the difference between the two ultrasound frequencies after adaptation to 200 Hz mechanical stimulation (*p* ≥ 0.196 in both cases).

Thus, this finding partially supported our hypothesis of frequency-specific adaptation, suggesting that a 30 s exposure to low-frequency mechanical adapting stimulus impairs the detection of low- but not high-frequency ultrasound test stimuli.

Following our preregistered analyses (https://osf.io/zwmnt), we then computed the percentage change in detection threshold from to the baseline for each condition (see Figure 2C). First, a series of one-sample t-tests comparing each condition against zero showed that all conditions except the 50 Hz mechanical – 200 Hz Ultrasound (t_11_ = 0.927; *p* = 0.374, Cohen’s d = 0.268) were significantly different from zero (i.e., baseline) (*p* ≤ 0.007 in all cases).

Next, a 2 (mechanical adapting stimulus: 50 Hz, 200 Hz) x 2 (ultrasound test stimulus: 50 Hz, 200 Hz) repeated measures ANOVA showed a marginally significant main effect of ultrasound frequency 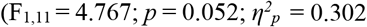.The main effect of mechanical frequency was not significant 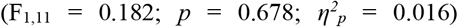. However, there was a significant interaction between factors 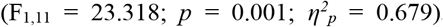. Two Bonferroni-corrected post-hoc comparisons showed that the 50 Hz mechanical adapting stimulus produced a significant increase in the detection threshold of an ultrasound test stimulus with a frequency of 50 Hz (54.16 % ± 37.30 %) but not 200 Hz (6.62 % ± 24.73 %; t_11_ = 3.474; *p* = 0.02; Cohen’s d = 0.790). In contrast, the 200 Hz mechanical adapting stimulus equally affected ultrasound test stimuli of both 50 Hz (28.83 % ± 30.30 %) and 200 Hz (37.33 % ± 15.91 %; t_11_ = −1.343; *p* = 0.824; Cohen’s d = −0.390; B_01_ = 1.677, error % = 0.005).

Thus, these results suggest that the relationship between mechanical adaptation and ultrasound detection is not explained by channel-specificity alone, but it also includes a directional asymmetric interaction between the different frequency-specific channels.

## III. EXPERIMENT 2

The results from Experiment 1 unveiled for the first time the impact of mechanical adaptation on the perception of ultrasound vibrotactile stimuli. However, previous studies on vibrotactile adaptation for mechanical stimuli have shown that the amplitude of the adapting stimulus is a crucial factor in determining the deleterious effect of adaptation on detection thresholds, with higher amplitudes inducing stronger adaptation effects [20], [21], [40], [41].

Yet, it remains unclear whether the effect of adapting amplitude can be similarly observed in channel-specific adaptation, particularly considering that our Experiment 1 showed only partial replication of the frequency-specific effect of adaptation.

Therefore, in Experiment 2, we used the same paradigm as in Experiment 1 to investigate the relationship between the amplitude of the adapting mechanical stimulus and the perception of the same ultrasound test stimuli used in Experiment 1. We hypothesized that stronger adapting stimuli would increase target detection thresholds (i.e., impair tactile perception) more than weaker adapting stimuli.

### A. METHODS

#### 1) PARTICIPANTS

The final sample size of Experiment 2 (n = 48; 37 female; mean age ± SD: 24 ± 4.1) was established a priori by means of a power analysis calculation on the results of our Experiment 1 (see second preregistration at https://osf.io/g8jft). To estimate the size of the effect of mechanical amplitude, we ran a 2 (mechanical frequency: 50, 200 Hz) x 2 (mechanical amplitude: 0, maximal amplitude possible) repeated measures ANOVA on the percentage change data from Experiment 1. The analysis showed that the main effect of amplitude 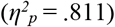 had a very large size, according to Cohen’s criteria [45]. Based on this effect size, an alpha level of 0.05, and a desired power level of 0.80, the projected sample size necessary to detect an effect of mechanical amplitude on ultrasound detection thresholds was determined to be n = 6. However, due to the addition of the new amplitude factor, Experiment 2 had a much larger number of conditions than Experiment 1 and hence had to be based on a between-participants design to avoid multiple, long testing sessions (> 5 h in total). Thus, to account for the methodological differences in Experiment 2, and for consistency with the previous experiment, we opted for a sample size of n = 12 for each condition deriving from our 2 (mechanical frequency: 50 Hz, 200 Hz) x 2 (ultrasound frequency: 50 Hz, 200 Hz) experimental design (see https://osf.io/g8jft).

A total of 57 healthy, right-handed participants were originally recruited for the study. However, based on preregistered exclusion criteria (see https://osf.io/g8jft), four participants from the M0-U50 condition and five participants from the M50-U50 condition were excluded after visual inspection of the data because they showed poor convergence (i.e., > 25% discrepancy) between ascending and descending staircases in the main task (see Procedure). The participants had not participated in Experiment 1.

All the methods and procedures employed in the study adhered to the Declaration of Helsinki and were approved by the UCL Research Ethics Committee. All participants were naïve regarding the hypotheses of the study, provided their written informed consent before the beginning of the experiment, and received a cash compensation of £9 per hour.

#### 2) EXPERIMENTAL DESIGN AND PROCEDURE

Experiment 2 had the same experimental setup as Experiment 1. However, detection thresholds for mid-air ultrasound stimuli were tested in a 2 (frequency of mechanical stimulus: 50 Hz, 200 Hz) x 2 (frequency of ultrasound stimulus: 50 Hz, 200 Hz) x 4 (amplitude of mechanical stimulus (0 N, 0.5 N, 1 N, 1.5 N for 200 Hz and 0 N, 0.08 N, 0.16 N, 0.24 N for 50 Hz) mixed design. The first two factors were tested between participants, while the mechanical amplitude factor was tested within participants.

This factor consisted of four equidistant levels of mechanical amplitude, ranging from no vibration at all (acting as a baseline for measuring ultrasound detection threshold in absence of mechanical adaptation) to the same amplitude tested in Experiment 1 (i.e., the highest amplitudes testable before incurring ceiling effects). Similar to Experiment 1, the amplitudes of the 50 Hz mechanical stimulus were set to the perceived isointensity of the and 200 Hz mechanical stimulus. To achieve this, we used the average isointensity values from Experiment 1 as the maximal amplitude of the 50 Hz mechanical stimulus, and then divided this value into four equidistant levels. This was done for the sake of reducing the duration of the testing session, and because the mechanical intensity matching data in Experiment 1 showed low variability, thus suggesting that using an average value should not generate high interindividual variability in the perception of the mechanical stimuli.

The detection threshold for ultrasound test stimuli were tested with the same psychophysical staircase procedure described in Experiment 1. Each participant performed a single testing session composed of four blocks (one for each mechanical amplitude level) in one of the four possible combinations of mechanical and ultrasound frequency conditions.

### C. RESULTS

The analysis of the participants’ raw detection thresholds showed similar statistical inference from the percentage change analysis (see Figure 3A and B). As in Experiment 1 and following our preregistration (see https://osf.io/g8jft) the main analyses were run on the percentage change between each condition and the 0 N, no adaptation baseline condition (see Figure 3C).

**FIGURE 3.**
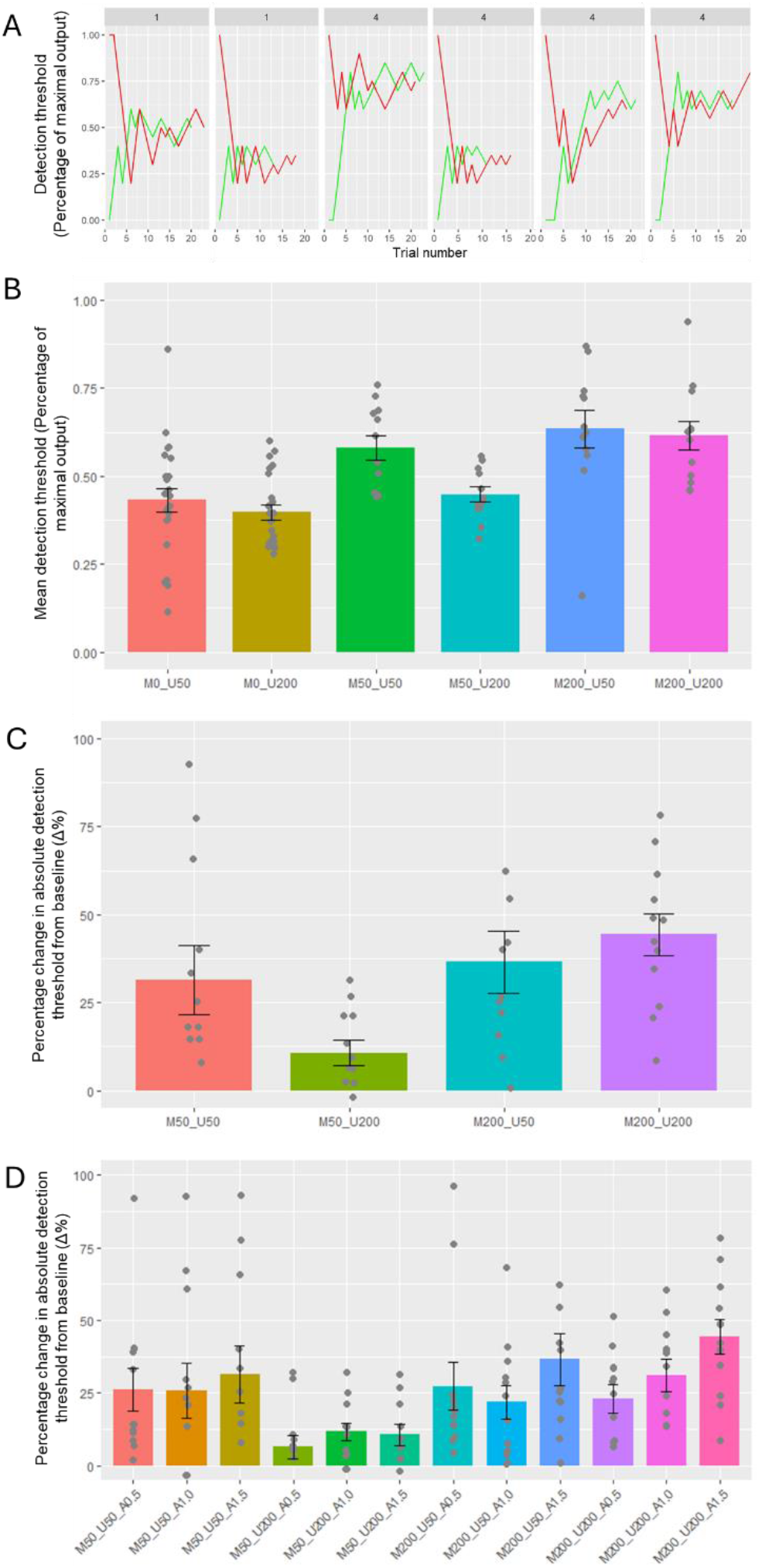
Results from Experiment 2. **A**. Staircase data from representative participants across conditions in Experiment 2 (P# 8: conditions M0_U50 and M50_U50; P# 1: conditions M0_U200 and M200_U200; P# 10: condition M50_U200; P# 16: condition M200_U50). Green and red lines show responses to ascending and descending staircases, respectively. **B**. Mean absolute detection thresholds for 50 Hz and 200 Hz ultrasound stimuli after the three mechanical adaptation conditions (no adaptation, 50 Hz, and 200 Hz). **C**. Percentage change in absolute detection thresholds from baseline for the conditions with 1.5 N amplitude replicated the results from Experiment 1. **D**. The main effect of amplitude of mechanical adapting stimulation was significant across conditions.

First, we tested whether the current experiment replicated the results from Experiment 1 by analysing the percentage change data from the conditions with an amplitude of 1.5 N, as in Experiment 1. A series of one-sample t-tests comparing each condition against zero showed that all adaptation conditions were significantly different from zero (i.e., baseline) (*p* ≤ 0.014 in all cases). Next, based on the results of Experiment 1, we ran two planned comparisons on the M50_U50 vs M50_U200 conditions and the M200_U50 vs M200_U200 conditions. The analyses showed that the 50 Hz mechanical adapting stimulus produced a significant increase in the detection threshold of an ultrasound test stimulus with a frequency of 50 Hz (31.49 % ± 33.83 %) but not 200 Hz (10.68 % ± 12.65 %; t_22_= 1.996; *p* = 0.026; Cohen’s d = 0.815). In contrast, the 200 Hz mechanical adapting stimulus equally affected ultrasound test stimuli of both low- (36.54 % ± 30.92 %) and high-frequencies (44.39 % ± 20.60; t_22_= −0.732; *p* = 0.380; Cohen’s d = −0.299; B_01_ = 2.203, error % = 0.017). Thus, these results confirmed Experiment 1 main result of asymmetric channel-specificity.

Finally, to test the effect of amplitude on vibrotactile adaptation, the percentage change data from all conditions were entered in a 2 (frequency of mechanical stimulus: 50 Hz, 200 Hz) x 2 (frequency of ultrasound stimulus: 50 Hz, 200 Hz) x 4 (amplitude of mechanical stimulus (0 N, 0.5 N, 1 N, 1.5 N for 200 Hz and 0 N, 0.08 N, 0.16 N, 0.24 N for 50 Hz) mixed ANOVA. The analysis showed a significant main effect of mechanical amplitude 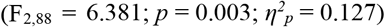. The finding was supported by a linear trend analysis showing a significant monotonic increase in ultrasound detection thresholds as a function of mechanical adaptation amplitude 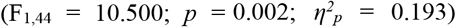, suggesting that the higher the amplitude of the mechanical adapting stimulus, the higher the increase in ultrasound detection thresholds. The main effect of frequency of mechanical stimulus was also significant 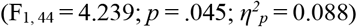, suggesting that, overall, the 50 Hz mechanical stimulation produced a significantly smaller increase in ultrasound detection thresholds (18.74 ± 24.53 %) compared to the 200 Hz mechanical stimulation (30.70 ± 23.82 %). The interaction between mechanical and ultrasound frequency was marginally significant 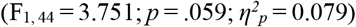, and it was explained by a larger difference between ultrasound thresholds after adaptation to a 50 Hz mechanical stimulus, but not after adaptation to a 200 Hz mechanical stimulus. Although the post-hoc comparison did not survive Bonferroni correction (*p* ≥ 0.062 in all cases), this trend was in line with our direct replication of Experiment 1, involving the strongest amplitudes only (see above). The main effect of frequency of ultrasound stimulation and all the other interactions were not statistically significant (*p* ≥ .102 in all cases).

Thus, overall, the main analyses of Experiment 2 confirmed our hypothesis that higher amplitudes of mechanical adapting stimuli produce a significantly stronger impairment of the detectability of ultrasound stimuli.

## IV. DISCUSSION

Mid-air ultrasound vibrotactile stimulation has been increasingly studied in recent years for its potential benefits in touchless communication and human-computer interaction. However, most real-world applications, including, e.g., alerting systems in cars, are charactherised by an inherent degree of environmental mechanical noise that could potentially impair the effectiveness of mid-air haptics. The current study provides the first systematic investigation of the effects of prolonged mechanical noise on perception of ultrasound mid-air stimuli of different frequencies.

We measured participants’ detection thresholds for ultrasound target stimuli before and after exposure to mechanical adapting stimuli of low and high frequencies and amplitudes. Across two experiments, including a preregistered replication, we found a strong effect of mechanical adaptation on ultrasound perception. Crucially, this effect was characterised by an interaction between the frequency of the mechanical and ultrasound stimulation, suggesting an asymmetrical frequency-specificity of adaptation. Finally, we also found that the increase in ultrasound detection thresholds induced by mechanical adaptation increased monotonically as a function of the amplitude of the adapting stimulus.

Our core finding of increased detection thresholds for mid-air ultrasound stimuli after mechanical adaptation is seemingly in contrast with the only previous study on the interaction between mechanical noise and ultrasound perception [39]. In that study, Špakov et al. showed that road vibration did not impair the driver’s ability to perceive the shape of different ultrasound stimuli [39]. However, the divergence between Špakov et al.’s results and ours can be likely explained by the multiple differences between their design and ours. First, Špakov et al. adopted a more naturalistic approach by using recorded or real vibrational noise from a moving car as a background mechanical stimulation delivered at the same time of the ultrasound stimulation. While the authors did not report the frequency and amplitude of their mechanical noise, the dominant fundamental frequencies of a moving vehicle are typically between 10 and 20 Hz [46], [47], [48], thus considerably lower than the mechanical frequencies tested in the present study. In contrast, our robotic setup allowed us to systematically manipulate both the frequency and the amplitude of the mechanical adapting stimulus and to design mechanical vibrations that were closely matched to the target ultrasound stimuli. Furthermore, while Špakov et al. tested participants’ ability to recognise mid-air shapes, we used standard psychophysical methods to measure the participants’ absolute detection thresholds for single-point ultrasound stimulation. Therefore, Špakov et al.’s study perhaps failed to detect adaptation because shape identification accuracy may be a less sensitive dependent variable than absolute detection thresholds.

Importantly, the present study also showed a significant interaction between the frequency of the mechanical adapting stimulus and the ultrasound target stimulus. In particular, we found that a brief (30 s) mechanical vibrotactile stimulation at 50 Hz selectively impaired detection of 50 Hz, but not 200 Hz ultrasound vibration. Conversely, mechanical vibrotactile stimulation at 200 Hz impaired the detection of both 50 Hz and 200 Hz ultrasound vibration in equal measure. Existing physiological evidence on mechanical-to-mechanical adaptation shows that adaptation to specific vibrotactile frequencies impairs perceptual thresholds of similar frequencies [14], [15], [23], [24], [25], [26]. This frequency-specific effect is commonly attributed to the desensitisation of different mechanoreceptive afferent channels [28], [29], [30], [31], [32], [33]. Yet, our results showed only partial frequency-specificity, suggesting that cross-adaptation between mechanical and ultrasound stimuli is asymmetrical and only occurs when the mechanical adapting stimulus has equal or higher frequency than the target ultrasound stimulus. This result can further explain the seeming difference between ours and Špakov et al.’s [39] results, suggesting that the lack of mechanical adaptation in their study might be due to the fact that their mechanical stimulation was considerably lower than the frequency of their ultrasound stimuli. This result of asymmetric adaptation is also in line with previous findings by Grosbois et al. [14] (see also [33], [49]), showing that the perception of a target 5 Hz stimulus was impaired by adaptation to a 5 Hz or 8 Hz stimulation, but not after a 2 Hz adapting stimulation. While the stimuli used by Grosbois et al. were about mechanical-to-mechanical adaptation and likely activated the same low-frequency afferent channel, our results extend this finding to the interaction between multiple mechanoreceptive channels and stimulation types.

Increasing evidence suggests that the PC channel also responds to low-frequency stimuli [50]. Therefore, a potential explanation for the asymmetric pattern of adaptation found in our study could be that both the PC channel and the RA channel were jointly activated by the 50 Hz target ultrasound stimulation. This could explain why the perception of both the 200 Hz and 50 Hz ultrasound target stimuli was equally impaired by the 200 Hz mechanical adapting stimulus, as in this case, the mechanoreceptor desensitisation of the PC channel would be shared across the two conditions.

Classical studies on mechanical-to-mechanical vibrotactile adaptation consistently show that the amplitude of the adapting stimulus is a key feature in determining the magnitude of the adaptation aftereffect [20], [21], [40], [41]. The results of our Experiment 2 confirmed that this is also the case for mechanical-to-ultrasound vibrotactile adaptation. We found a positive linear relationship between the participants’ ultrasound detection thresholds as a function of the amplitude of the mechanical adapting stimulus, with higher amplitudes producing higher detection thresholds.

While these results show that mechanical adaptation could be a serious concern for the effectiveness of mid-air ultrasound haptics, they also have important implications in terms of potential solutions for real-world applications. In particular, the asymmetrical aftereffect we found across our two experiments suggests that the effect of mechanical adaptation on ultrasound perception only takes place when the adapting stimulus has equal or higher frequency than the target stimulus. Therefore, to minimise adaptation, applications involving mid-air haptics in noisy environment could be equipped with sensors that constantly monitor the background mechanical noise in real-time and ensure that the frequency of the mid-air stimulus is consistently higher than the detected mechanical vibration noise. Similarly, our results on the linear relationship between mechanical amplitude and ultrasound detection suggest that mid-air haptics might be ineffective when the amplitude of the mechanical noise is too high. Therefore, real-world applications might include a closed-loop system that modulates the amplitude of the ultrasound stimulation as a function of the frequency and amplitude of the mechanical environmental noise.

In our experiments, both adapting and target vibrotactile stimuli were delivered to the palm of the participants’ hand only. This was done so to mimic a scenario where a driver holds the steering wheel of a vehicle right before interacting with a hypothetical mid-air haptics infotainment system. However, environmental mechanical noise in a moving vehicle is likely to apply to the whole body. Thus, future studies should investigate the potential effect of mechanical adaptation on ultrasound sensitivity of multiple or distant body parts.

## ACKNOWLEDGMENT

This study was supported by the TOUCHLESS European Union’s Horizon 2020 research and innovation programme under grant agreement No 101017746 and by a Behavioural Insights Exchange programme between UCL and Ultraleap. W.F. was also supported by a UKRI Future Leader Fellowship (grand number: MR/W013476/1. We thank Katherine Wang for helping with data collection.

## AUTHOR CONTRIBUTIONS

Conceptualization: A.C., W.F., and P.H.; methodology: A.C., T.H., W.F., and P.H.; software: A.C., T.H., and W.F.; formal analysis and visualization: A.C.; validation and investigation: T.H. and A.C.; resources: P.H. and W.F.; data curation: T.H. and A.C.; writing – original draft: A.C. and T.H.; writing – review & editing: A.C., W.F., and P.H.; supervision, project administration, and funding acquisition: P.H..

